# Microbial diversity and ecological complexity emerging from environmental variation and horizontal gene transfer in a simple mathematical model

**DOI:** 10.1101/2024.01.17.576128

**Authors:** Sanasar G. Babajanyan, Sofya K. Garushyants, Yuri I. Wolf, Eugene V. Koonin

**Affiliations:** National Center for Biotechnology Information, National Library of Medicine, National Institutes of Health, Bethesda, MD 20894, USA

## Abstract

Microbiomes are generally characterized by high diversity of coexisting microbial species and strains that remains stable within a broad range of conditions. However, under fixed conditions, microbial ecology conforms with the exclusion principle under which two populations competing for the same resource within the same niche cannot coexist because the less fit population inevitably goes extinct. To explore the conditions for stabilization of microbial diversity, we developed a simple mathematical model consisting of two competing populations that could exchange a single gene allele via horizontal gene transfer (HGT). We found that, although in a fixed environment, with unbiased HGT, the system obeyed the exclusion principle, in an oscillating environment, within large regions of the phase space bounded by the rates of reproduction and HGT, the two populations coexist. Moreover, depending on the parameter combination, all three major types of symbiosis obtained, namely, pure competition, host-parasite relationship and mutualism. In each of these regimes, certain parameter combinations provided for synergy, that is, a greater total abundance of both populations compared to the abundance of the winning population in the fixed environments. These findings show that basic phenomena that are universal in microbial communities, environmental variation and HGT, provide for stabilization of microbial diversity and ecological complexity.

## BACKGROUND

The extensive recent efforts in metagenomics have led to the realization that microbiomes, for example, the well-studied gut microbial communities, are surprisingly stable on the species level over long periods of time [1–3]. Changes in microbiome content caused by such factors as antibiotics intervention or diet change can lead to obesity, inflammatory bowel disease and other pathological conditions, so that microbiome stability is essential for human health [4]. Bacterial and archaeal species within the microbiome have metabolic networks that complement each other, and are thought to cooperate [5–7]. However, there are also indications of strong interspecies competition in complex microbial communities such as soil [8].

With the development of single cell metagenomics, finer structure of microbiomes was discovered, demonstrating that most of the constituent bacterial species are represented by multiple, closely related strains [9–12]. The coexistence of multiple strains has been shown to be common, and moreover, strain composition tends to be stable over extended periods of time [10, 13, 14]. It is generally assumed that the observed stability of the strain composition can be accounted for by models with no competition between strains [15]. Whereas prevalence of cooperation has been demonstrated for different microbial species, metabolic networks of strains of the same species are closely similar, so that competition for resources is expected to occur. Indeed, attempts on administration of probiotics have shown that bacteria, occupying the same niche as the species that is already dominant in the microbiome, are eliminated [16, 17].

A key process in the evolution of prokaryotes is the extensive horizontal gene transfer (HGT) that occurs via transformation, transduction or conjugation (reviewed in [18]). The size of the transferred DNA fragments varies from several hundred nucleotides to segments of many kilobases encompassing multiple genes. Horizontal gene transfer is thought to be essential for the survival of microbial populations [19] and appears to be largely responsible for rapid adaptation to new environments and even for the emergence of major clades of bacteria and archaea with distinct lifestyles, such as acetoclastic methanogens or extreme halophiles [20, 21]. The rate of HGT is highly non-uniform across the tree of life and across environments [22]. Indeed, there is growing evidence that HGT rate can be influenced by the environmental changes. For example, it has been shown that natural competence of the bacterium Staphylococcus aureus is induced by reactive oxygen species, antibiotics and host defenses [23, 24]. Importantly, the HGT rate is much higher between closely related organisms compared to distantly related ones thanks, primarily, to the high rate of homologous recombination [25–27].

In recent years, microbiome stability and variation have become a major problem in microbiology, with important implications for human health [28, 29]. Horizontal gene transfer plays a crucial role in the preservation of genome diversity [30–36] but the effects of environmental variations on the microbiome composition remain poorly understood. Evolutionary processes in populations in time-varying environments can drastically differ from those in fixed environments [37–48]. The complexity of microbial communities including multiple strains that persist for extended periods of time and widespread HGT among them call for developing theoretical models describing the relationships between strains within complex microbial communities, that are expected to compete with each other, but also constantly exchange genes via HGT.

Here we describe the simplest possible model that includes competition and HGT between two microbial populations differing by a single gene allele in a fixed or oscillating environment. The time-varying environment drastically changes the evolutionary scenarios observed in a fixed environment. We show that in this setting environmental variation plays a crucial role in the preservation of genome diversity. The coarse-graining modeling presented here allows us to uncover different types of symbiotic relationships that arise due to environmental variation, spanning all possible scenarios, from pure competition to mutualism [49–52].

## MODEL OF MICROBIAL POPULATIONS DYNAMICS

We explore a dynamic evolutionary scenario where two populations engage in both within- and between-population competition for shared resources. The two populations differ by a single gene allele, resulting in differential fitness. HGT plays a key role in the model, mediating the exchange of genetic material between the two populations by copying the gene from the donor and replacing the corresponding gene in the recipient. For instance, consider the case when population *A* harbors allele 1, and population *B* carries allele −1. If *A* is the donor and *B* is the recipient, then the allele from *A* replaces that in *B*, and as a result, the recipient changes its type *B* → *A*, and the population sizes of *A* and *B* increase and decrease by one, respectively. The donor and recipient entities are selected randomly within their respective populations. The rates of gene transfer vary depending on whether A or B is the donor. Inactivation of genes by mutations is disregarded.

To capture the dynamics of this HGT-mediated interaction mathematically, we derive a system of differential equations that describes the continuous variation of population sizes over time:

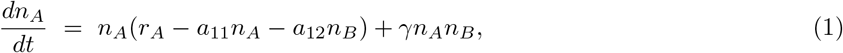

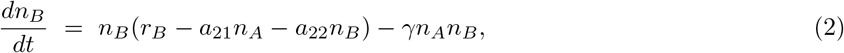

where *n*_*A*_(*t*) and *n*_*B*_(*t*) are population sizes of *A* and *B* at time *t*, respectively. *r*_*A*_ *>* 0 and *r*_*B*_ *>* 0 are reproduction rates of *A* and *B*, respectively. *a*_*ii*_ and *a*_*ij*_ *i ≠ j* = 1, 2 are within- and between- population competition rates. *γ* is the balance of gene transfer between *A* and *B* types, i.e *γ >* 0 corresponds to the positive gene flow from *B* to *A*, when the HGT rate from *B* to *A* is greater than the HGT rate from *A* to *B*. The rates describing the competition, both within and between two populations, are assumed to be positive *a*_*ij*_ *>* 0, *i, j* = 1, 2, and the fitness of each population is a decreasing function of the presence of competing population, corresponding to a purely competitive ecosystem in the absence of HGT. The detailed derivation of these equations from elementary processes, based on previous work [53, 54], is provided in the Additional File1. For the purpose of the present manuscript, we will describe the system of (1, 2) by the equivalent representation through the total abundance of both populations *N* = *n*_*A*_ + *n*_*B*_ and fractions of each population *p*_*A*_ = *n*_*A*_*/N, p*_*B*_ = *n*_*B*_*/N*, such that *p*_*A*_ + *p*_*B*_ = 1. From (1,2), we obtain the following dynamical system representing the time variations of total abundance of both populations and the fraction of each population:

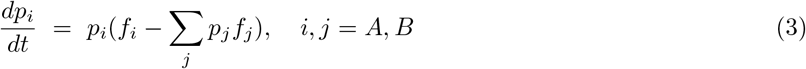

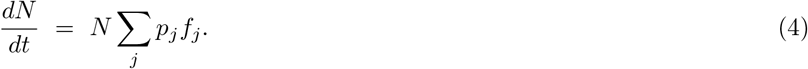

Equation (3) represents well-known replicator dynamics in evolutionary game theory [55–59]. However, the crucial difference between (3,4) and standard replicator dynamics is the dependence of fitness functions *f*_*i*_ on the total population size *N*. The fitness of each population can be represented as the expected payoff obtained from a 2 *×* 2 symmetric game with the payoff matrix shown in Fig.1. The payoffs of each strategy/population *A* and *B* depend on the total abundance *N* of both populations. The time variation of the total abundance of both populations is defined by the mean fitness of both populations. In the equilibrium states, when the right-hand sides of both equations (3,4) nullify, the mean fitness and the fitness of each population represented in the equilibrium state nullify as well. Solving the right hand-side of (3,4) with respect to the fractions of each population (taking into an account the normalization *p*_*A*_ + *p*_*B*_ = 1) and total population abundance, we find 4 possible non-trivial outcomes: 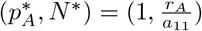 – extinction of population *B*, 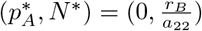 – extinction of population _*A*,_ 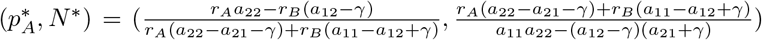_– if_ 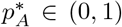, along with *N*^*^ *>* 0, then depending on the stability of equilibrium the outcome is either coexistence of both populations or bistability, where extinction of a population depends on the initial condition, and the interior unstable equilibrium *p*^*^ separates basins of attraction of two stable equilibrium states representing extinction of either *A* or *B* populations. The trivial equilibrium state, corresponding to the extinction of both populations, is always unstable for positive reproduction rates *r*_*i*_ *>* 0, *i* = *A, B*.

**FIG. 1:**
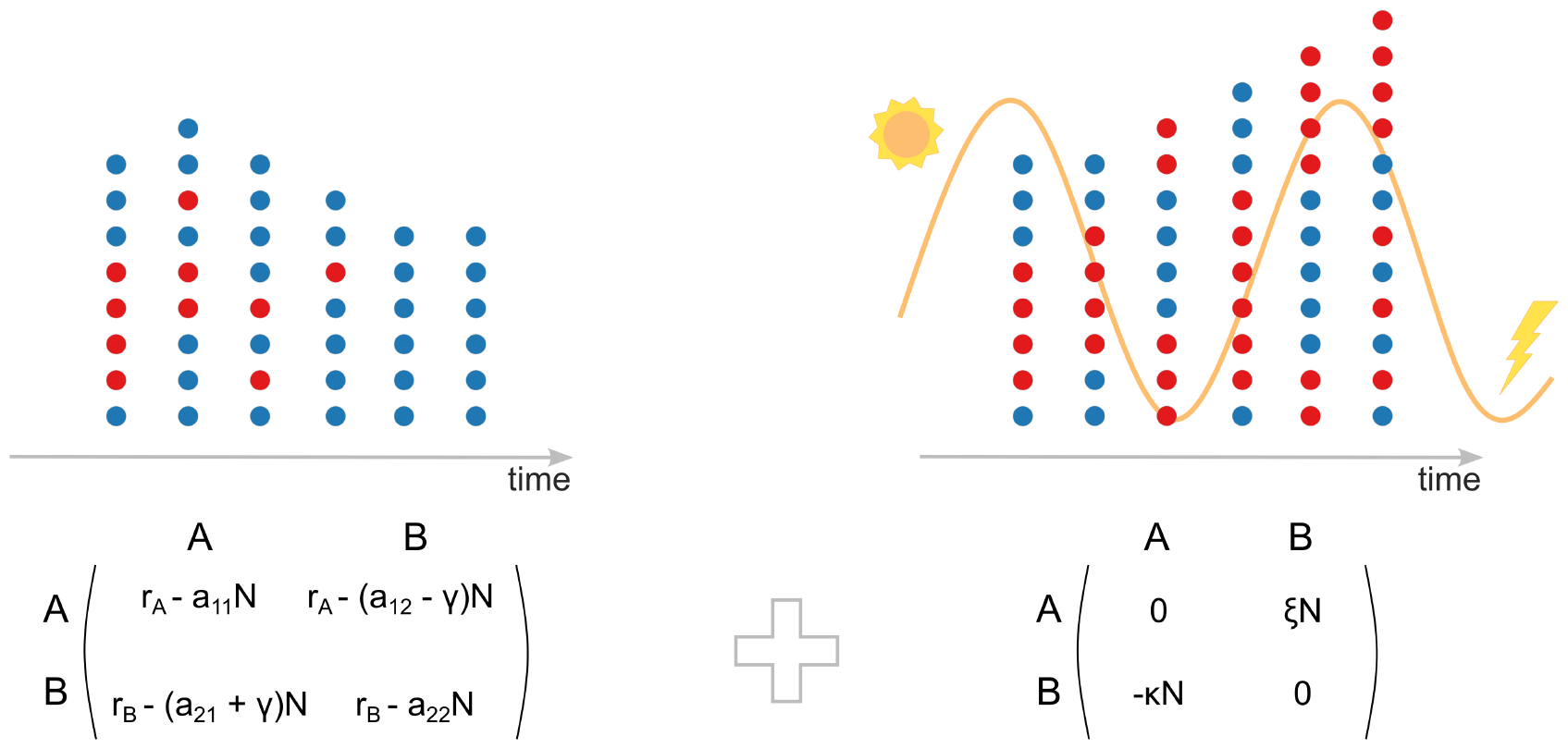
Evolutionary dynamics of two competing populations with HGT in a fixed and in a fast-oscillating environment. Left, fixed/averaged environment; right, oscillating environment. The coarse-grained time variations of the fractions of each population and the total abundance of both populations are described by the replicator dynamics with density dependent payoff matrices. The fitness of each population is comprised of two terms obtained from two different evolutionary games. These evolutionary games represent the interaction between the two populations in the fixed/averaged and varying environments (left and right matrices, respectively). In the left matrix, *r*_*A*_ and *r*_*B*_ are reproduction rates of the populations *A* and *B* in the averaged/fixed environment, respectively. The competitions within and between the two populations are given by *a*_*ij*_, *i, j* = 1, 2 rates. *γ* is the gene transfer balance in the averaged/fixed environment. *ξ* and *κ* represent environmental variations in the model, these are defined by the product of oscillatory parts of reproduction and gene transfer balance rates, defined in (11). *N* is the total abundance of both populations.

The stability of equilibrium states with the trivial composition of the total population is defined by the following conditions:

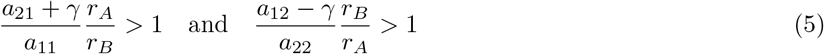

for 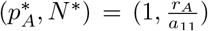 and 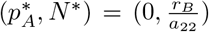, respectively. For *γ* = 0, the conditions (5) link between-populations and within-populations competition with the balance of the reproduction rates. Indeed for *r*_*A*_ = *r*_*B*_, a given population outcompetes the opponent if the competition within the population of the winner is less severe than the competition between-populations.

If both conditions in (5) are satisfied simultaneously, along with 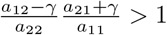, then one obtains a bistable dynamical system, where the outcome of competition depends on both the initial abundance and fractions of both populations. The last condition is linked to the sign of the determinant of the community matrix, represented via the competition and gene transfer balance rates

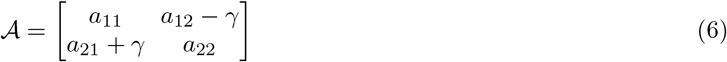

Coexistence is obtained, once both conditions in (5) are violated simultaneously along with

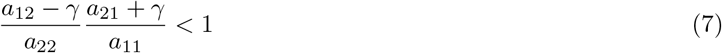

that is the Det 𝒜 *>* 0.

So far, we analyzed the model for general competition and reproduction rates, without imposing any other condition on the rates except for being positive. Therefore, our initial assumption is that populations *A* and *B* differ only by a single gene allele, and we further assume that the alleles of this gene affect only the reproduction rates *r*_*A*_ and *r*_*B*_, whereas the rates describing the competition within and between populations are independent of the gene allele variation, and are equal, *a*_*ij*_ = *a*. Thus, for equal reproduction rates *r*_*A*_ = *r*_*B*_ = *r* and balanced gene transfer between two populations *γ* = 0, the total abundance of both populations is always equal to 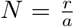. Coexistence of both populations could not be achieved in the case of balanced gene transfer *γ* = 0 and unequal reproduction rates *r*_*A*_ *≠ r*_*B*_, inasmuch as it is impossible to violate both conditions in (5) simultaneously. This impossibility of coexistence between populations corresponds to the exclusion principle which states that, if both populations compete for the same resources, then, one of them will be eliminated, under equal competition rates *a*_*ij*_ = *a*; in our setting, the exclusion principle applies in the case of balanced gene transfer *γ* = 0 [60–62].

In the case of unbalanced gene transfer (*γ≠* 0) coexistence of the two competing populations is possible if

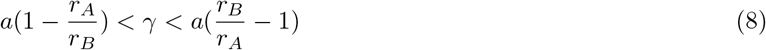

that is, both conditions in (5) are violated and (7) is satisfied simultaneously, under equal competition rates *a*_*ij*_ = *a*.

The above conclusions were obtained for a fixed environment. We now consider the same evolutionary process in a fluctuating environment. We assume that the environment oscillates faster than the characteristic timescale of the population dynamics, that is, both populations experience many changes of the environment before reaching the dynamic equilibrium. The environmental variations are incorporated in the model by assuming that the reproduction rates and the gene transfer balance rate are periodical function *r*_*i*_(*τ* +2*π*) = *r*_*i*_(*τ*), *i* = *A, B* and *γ*(*τ* + 2*π*) = *γ*(*τ*) with period 2*π/ω*, where *τ* = *ωt, ω >* 1 describes fast variations of the environment. Here, *t* is the slow-time, that represents coarse-grained variations in competing populations. The coarse-graining procedure is based on the elimination of the fast-oscillating terms by identifying the dynamical impact of these terms on the slow-time behavior. That is, the fast-varying terms introduce an “effective potential” in the slow-time behavior, and these fast-oscillating terms are themselves affected by the quantities varying in the slow-time. The time dependent reproduction and gene transfer balance rates are represented as the sum of fixed and oscillatory parts 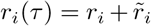 and 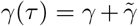, where the time averages of oscillatory parts nullify over a period of environmental variations, that is 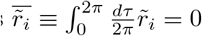 and 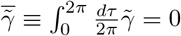. That is, *r*_*i*_ and *γ* represent the reproduction and gene transfer balance rates, respectively, in the averaged/fixed environment. Considering the evolutionary processes with averaged rates yields the system (3, 4) and the conclusions obtained for the fixed environment.

The derivation of the slow-time behavior is given in the Additional File1; here, we briefly outline the assumptions and implementation steps of this derivation. We seek the solution for the composition and total abundance of both populations as a sum of slow varying and oscillating parts, that is *p*_*A*_ = *p*_*A*_(*t*) + *π*_*A*_(*p*_*A*_(*t*), *N* (*t*), *τ*) and *N* = *N* (*t*) + 𝒱(*p*_*A*_(*t*), *N* (*t*), *τ*), respectively. The fast oscillating parts yield zero average over the period of environmental variations 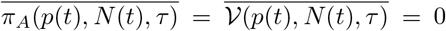. Note that the fast oscillating quantities *π*_*A*_(*p*(*t*), *N* (*t*), *τ*) and *V*(*p*(*t*), *N* (*t*), *τ*) depend on the slow varying counterparts *p*(*t*) and *N* (*t*). Putting the expressions of time-dependent rates *r*(*τ*), *γ*(*τ*) together with *p*_*A*_ and *N* back into (3,4), expanding the right hand side over *π*_*A*_, *𝒱*, seeking the fast oscillating parts as a power series with 1*/ω* as the order parameter, that is 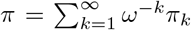 and 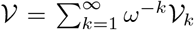, separating the quantities varying in different time scales, averaging over a period of environmental variations and keeping the terms up to *O*(1*/ω*^2^), we find the slow time behavior of *p*_*A*_(*t*) and *N* (*t*)

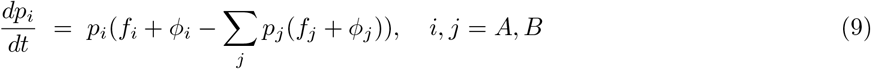

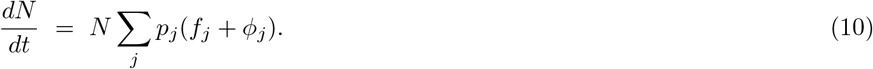

Thus, in the oscillating environment, the coarse-grained variations of the fraction of each population and total abundance of both populations, again, are given by the replicator dynamics, but the fitness of each population obtains an additional term due to the oscillating environment (Fig.1). Note, that *f*_*i*_, *i* = *A, B* is given by the average/fixed rates, as in (3,4). The new terms *ϕ*_*i*_ are expressed through the oscillating rates of reproduction and gene transfer balance as follows

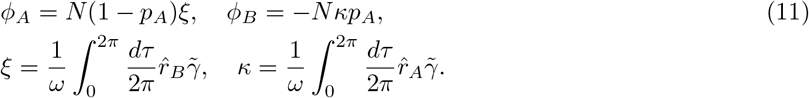

In (11), 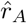 and 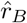 denote the primitives of 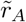 and 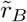, respectively, that is 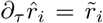 The existence and stability of possible equilibria of (9,10) are described by the conditions (5,7), where the between-population competition rates are substituted as follows *a*_12_ → *a*_12_ − *ξ* and *a*_21_ → *a*_21_ + *κ*. The initial conditions for (9,10) are found by solving the following system of algebraic equations 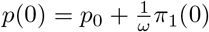 and 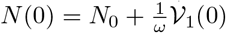, where *p*_0_, *N*_0_ are initial conditions for (9,10), *p*(0), *N* (0) are the initial conditions for (3,4) with time-dependent payoffs and *π*_1_(0), *𝒱*_1_(0) are the leading order oscillating terms proportional to *O*(1*/ω*). The latter terms are represented through *p*_0_, *N*_0_ and time-dependent rates 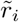 and 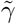 (see Supplementary Information for further details).

## RESULTS

### Environmental oscillations enable coexistence of competing populations

In the present model, oscillations in the environment induce a new evolutionary game between the competing populations (Fig.1). The emergence of the new game results from the coarse-graining of the effects of environmental variations on the competing populations. Specifically, the fast-oscillating terms induce an “effective potential”, a new game that alters the slow-time dynamics of the fractions of the two populations and total population abundance of both populations. The coarse-grained variations of the total population structure and abundance are defined by additive fitnesses obtained from two distinct games, one of which represents the evolutionary process in the fixed/averaged environment, and the other, emerging game represents the coarse-grained impact of the oscillating environment (11). The fitness of each population in the new game is expressed through the overlap in the oscillations of gene transfer balance rate 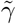 and the reproduction rate of the competitor over the period of environmental variations. The impact of the environmental variations is trivial if the oscillating part of the reproduction rates and gene transfer rate is the same, that is if *r*_*i*_(*τ*), *γ*(*τ*) ∝ *g*(*τ*), then *ξ, κ* = 0. To substantially influence population growth and composition compared to the fixed environment, environmental variations must differentially affect the reproduction and gene transfer balance rates. The new payoffs *ξ, κ* nullify when *ω* → ∞ for both the bounded oscillations of reproduction and gene transfer rates, as it follows from (11). That is, if environmental oscillations are too fast, then only the averages of reproduction and gene transfer balance determine the evolutionary outcome.

The payoff structure of the emerged game is simple for any positive population abundance *N*. These payoffs are fixed once the time dependence of reproduction and gene transfer balance rate is known. Then, either there is a dominant strategy in the game (that is, either *A* or *B* always win in the new game)–if *ξ* and *κ* have the same sign (for example, *A* dominates *B* if *ξ, κ >* 0), or there is a non-trivial equilibrium for any *N* if *ξ* and *κ* have different signs [56, 58, 63].

Environmental oscillation can result in coexistence of the two populations in a regime for which it is impossible in a fixed environment, that is, when gene transfer is balanced (*γ* = 0) between competing populations and the allele difference affects only the reproduction rates (*a*_*ij*_ = *a*). In a fixed/averaged environment, the exclusion principle applies so that only the population with the higher reproduction rate survives, giving the total abundance of both populations as 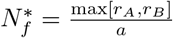.

Assuming that *a*_*ij*_ = *a* and gene transfer is balanced on average (*γ* = 0), we substitute *a*_12_ → *a*−*ξ, a*_21_ → *a*+*κ* in (5,7) and find the following conditions for the existence and stability of the coexistence of the two populations in the coarse-grained dynamics (9,10)

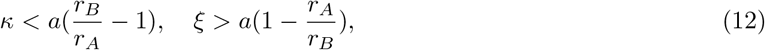

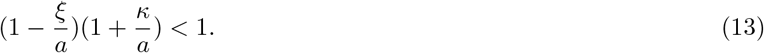

Let us assume that the average reproduction rates of both populations are almost equal, 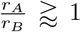. Then, from (12), it follows that, in a varying environment coexistence is possible for *ξ >* 0 and *κ <* 0, whereas, from (5), it follows that in a fixed environment coexistence is impossible if *a*_*ij*_ = *a* and *γ* = 0. From (12) it follows that environmental variations dampen the between-population competition such that the competition within a population becomes more severe than the competition between the populations. This is the case because environmental variation shifts the between-population competition rates *a*_12_ = *a* − *ξ < a* and *a*_21_ = *a* + *κ < a*, as follows from emerged fitness terms (11), thus reducing between-population competition compared to the within-population competition.

The total abundance of both populations in the coexisting equilibrium state, obtained from the coarse-grained representation of the oscillating environment, is

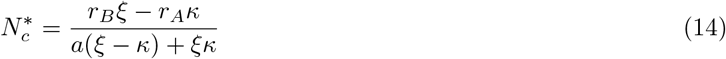

The conditions (12) and (13) provide 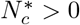, ensuring that the numerator and denominator are both positive,respectively.

### The phase space of interactions between populations

Environmental oscillations can result not only in coexistence but also in a synergistic interaction between the two competing populations for given reproduction rates *r*_*A*_ and *r*_*B*_. By synergy, we refer to the situations when the presence of the two populations in the equilibrium state increases the total abundance of both populations compared to the total abundance in the equilibrium state in a fixed environment 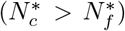. The synergy emerges, because the presence of both populations increases the utilization of available resources in the environment. Comparing, the total abundance of both populations in the varying environment-induced coexistence regime (14) and that in the fixed/averaged environment 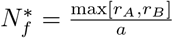, we find that

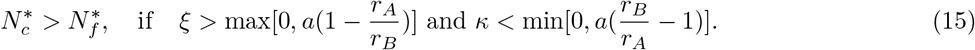

Assuming *r*_*A*_ *> r*_*B*_, we now consider different regions of the parameter space bounded by *ξ* and *κ*, where (12) is satisfied, corresponding to the different scenarios. The separation of these regions must be complemented with (13) for the coexistence equilibrium to be stable and the total abundance of the two populations to remain positive.

#### Coexistence without synergy: pure competition and parasitism

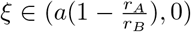 and 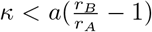, region I in Fig.2a.

**FIG. 2:**
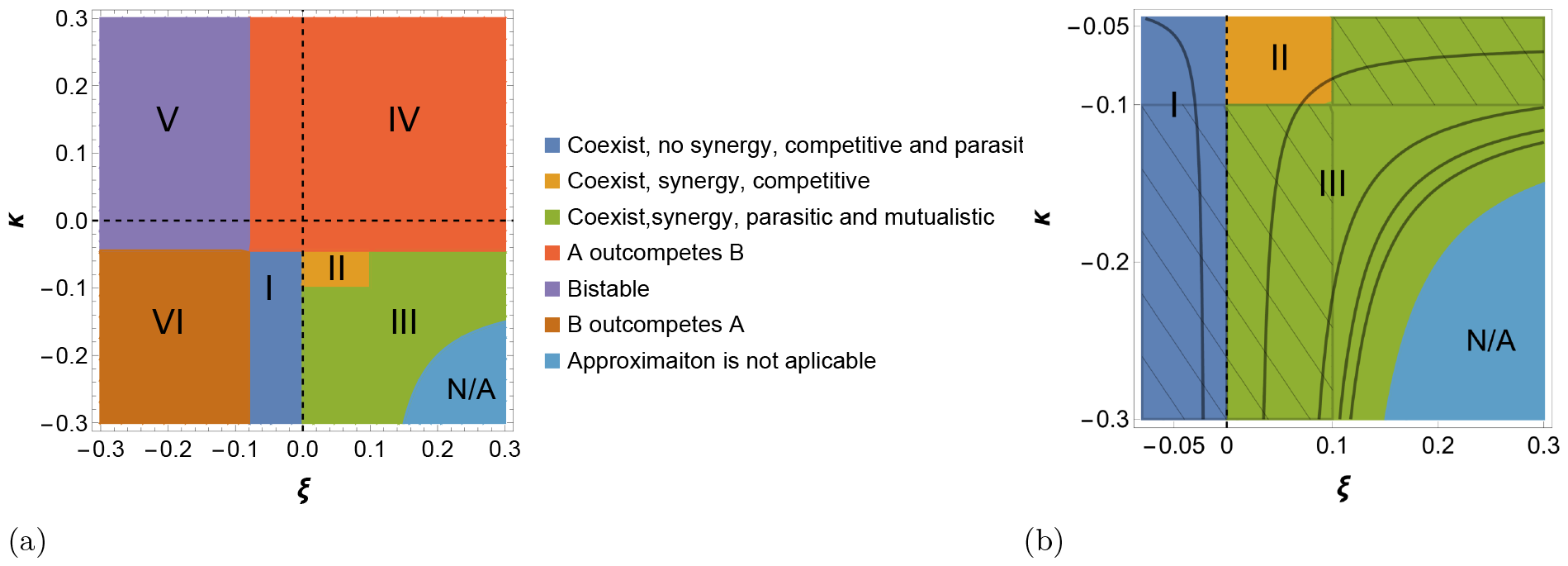
Phase space for possible outcomes of the coarse-grained description of two competing populations in an oscillating environment (9,10), for different values of environment-induced fitness terms. (a), partitioning of the phase space into distinct regions (see color code). Regions I, II, and III correspond to the coexistence of the two populations. In the regions II and III, there is synergy between the competing populations, that is, in the unique equilibrium state, the total abundance of both populations is greater than that in the fixed environment (for the considered values of the model parameters, *A* outcompetes *B* in the fixed environment). In regions IV, *A* outcompetes *B*, like in a fixed environment, V is the region of bistability, where either *A* or *B* wins depending on the initial state, and in region VI, *B* outcompetes *A*, opposite to the outcome in the fixed environment. Regions IV, V and VI are defined by violations of one or more of the conditions (12,13). *N/A* represents the region where the coarse-grained approach fails, that is (12) holds, but (13) is violated. (b), detailed structure of the regions I, II, III and N*/*A. The hatched areas represent the regions where the fitnesses of the two populations are differently affected by the presence of the competitor. The solid black curves show the total abundance of both population defined by (14), increasing from I to III. The model parameters are *r*_*A*_ = 1.8, *r*_*B*_ = 1, *γ* = 0 and *a* = 0.1.

In this region, the fitness of *A*, the winner in the fixed environment, is lower than it would be in the latter case. Indeed, the total fitness of *A* is *f*_*A*_ + *ϕ*_*A*_, where *ϕ*_*A*_ = *N* (1 − *p*_*A*_)*ξ <* 0 (Fig.1). Conversely, the population *B* obtains a positive fitness surplus due to the environmental oscillations *ϕ*_*B*_ = −*Np*_*A*_*κ >* 0. In the emerged game, *B* always has an advantage over *A*. Thus, *A* dominates over *B* in a fixed environment, but *B* has the advantage in the new game induced by the oscillating environment. In this region, there is coexistence but no synergy between two populations, that is, the total abundance of the two populations is lower than that in the fixed environment with same parameters, 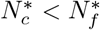.

The coarse-grained approach allows us to identify distinct sub-regions of different symbiotic relations between the two populations, that is, define the impact of the opponent on the fitness of a population. Region I includes two sub-regions that correspond, respectively, to competition and parasitism. In the competitive sub-region 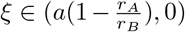 and 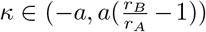, the fitness of each population is a decreasing function on the fraction of the competitor, that is 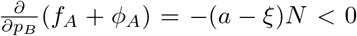 and 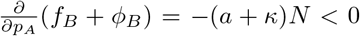. The latter quantity changes its sign for *κ <* −*a*, so in the corresponding sub-region, *B* is a parasite of *A*, dashed sub-region of I in Fig.2b. Due to environmental oscillations, the above derivatives can change their signs in the case of time-dependent rates, hence, the type of the symbiotic relation between the two populations cannot be inferred without coarse-graining.

#### Synergy between populations: pure competition

*ξ* ∈ (0, *a*) and 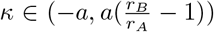, region II in Fig.2a. The payoffs of the emerged game are positive *Nξ*, −*Nκ >* 0, in contrast to region I. Thus, the two populations coexist due to the balance between the two games. The fitness of each population is a decreasing function of the competitor’s fraction 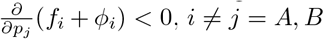, *i≠ j* = *A, B*, hence, the interaction is purely competitive. Nevertheless, the total abundance of both populations is greater than that in the fixed environment 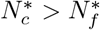, as follows from (15), corresponding to increased utilization of the available resources.

#### Synergy: parasitism and mutualism

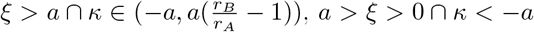, *a > ξ >* 0 ∩ *κ <* −*a*, and *ξ > a* ∩ *κ <* −*a*, corresponding to parasitism and mutualism, respectively, in two sub-regions of region III (Fig.2a,2b).

In the sub-region 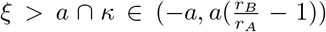, the fitness of *A* is positively affected by the presence of the competitor *B*, that is 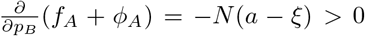, whereas the fitness of *B* is negatively impacted by *A*, that is 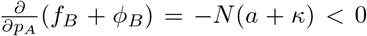, so that *A* is a parasite of *B*. Conversely, in the sub-region, *a > ξ >* 0 ∩ *κ <* −*a*, fitness of *A* is negatively impacted by *B*, whereas *A* positively affects the fitness of *B*. Accordingly, the host-parasite relationship is reversed.

The symbiosis between the two populations is mutualistic when *ξ > a* ∩ *κ <* −*a*. In this sub-region of region III, the fitness of each population is an increasing function of the fraction of the other population 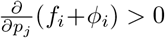, *i≠ j* = *A, B* (2b). The total abundance of both populations is greater than that in the fixed environment in both parasitic and mutualistic symbiosis, that is, both types of symbiosis are synergistic.

The total abundance of both populations (14), 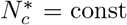, increases from region I to region III. The total abundance of both populations in the fixed environment corresponds to *ξ* = 0 line, thus, by substituting *ξ* = 0 in (14), one recovers 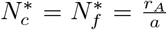. In the fixed environment, only population *A* survives, so that the total abundance equals to the abundance of *A*, whereas in the oscillating environment, both *A* and *B* are present in equilibrium with fractions 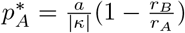 and 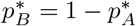, respectively.

The coarse-grained approach fails as one gets closer to the curve *a*(*ξ* − *κ*) + *ξκ* = 0, where 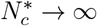, whereas the time-dependent solution is bounded (N*/*A regions in Fig.2). Unbounded growth corresponds to violation of (13) that holds as equality. Below the curve defined by *a*(*ξ* − *κ*) + *ξκ* = 0, 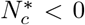. The fraction of each population satisfy 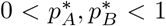. Thus, there is a non-trivial composition of both populations, but, there is no physical level of total abundance. This is one of the differences from the well-known replicator dynamics, which deals with the composition of the total population only. The reason behind the failure is the unbounded growth of both populations (see Supplementary Information for details on the limitations of the coarse-grained approach).

Figure 3 illustrates the solutions for the fractions and total abundance of two populations, for both time-dependent rates and the coarse-grained behavior (9,10).

**FIG. 3:**
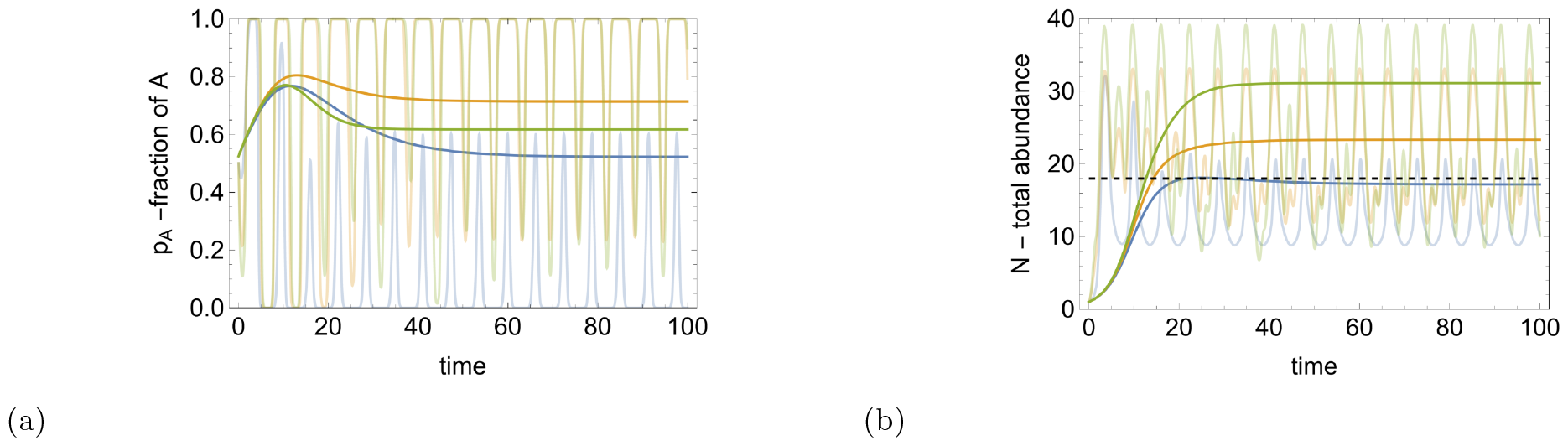
Fractions and total abundance of the two populations. (a), fraction of the population *A* for different values of *ξ* and *κ*. (b), total abundance of both populations for different values of *ξ* and *κ*. Oscillating curves show the solutions of (3,4) with time-dependent reproduction and gene transfer balance rates. (*ξ, κ*) = (−0.01, −0.08) (region I in Fig.2; blue curve), (*ξ, κ*) = (0.08, −0.08) (region II; orange curve) and (*ξ, κ*) = (0.11, −0.11) (region III; green curve). The time dependent rates are as follows 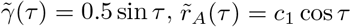 and 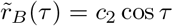, where (*c*_1_, *c*_2_) = (−1.6, −0.2), (−1.6, 1.6) and (−2.2, 2.2) for the blue, orange and green curves, respectively. The remaining model parameters are *a* = 0.1, *γ* = 0, *ω* = 5, *r*_*A*_ = 1.8 and *r*_*B*_ = 1. The dashed line in (b) represents the total abundance of both populations in the fixed environment 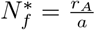.

The ordering of fractions and total abundance does not follow a uniform pattern across the different scenarios of interaction between the populations. In the case of pure competition with synergy, the equilibrium fraction of population *A* surpasses that in the mutualistic scenario. However, the total abundance of both populations is greater in the mutualistic scenario than in the pure competitive case. The scenario with no synergy (region I in Fig.2) yields the smallest total abundance for both populations as well as, for the fraction of population *A*. These observations imply complex relationships between competition, synergy, and mutualistic symbiosis, which affect both population fractions and overall abundance.

So far, we addressed the emergence of coexistence between two competing populations due to fast environmental variations. In other parts of the phase space, however, the environmental variations can preserve the outcome of the competition in the fixed environment, that is, *A* outcompetes *B* (region IV in Fig.2), or reverse it (region VI) resulting in the domination of *B* and extinction of *A*. The former scenario occurs if 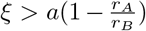 and 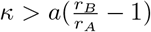, that is, the inequality (12) is reversed for *κ*. Conversely, *B* outcompetes *A* if the inequality is reversed for *ξ*, whereas that for *κ* holds. Finally, the bistable dynamics (region V) is observed if (12) and (13) are simultaneously violated.

Unbalanced gene transfer (*γ≠* 0) in a fixed environment can result in stable coexistence of the two populations (8). In this case, environmental variations can alter the composition and the total abundance of the two populations and even destroy the coexistence of the two competing populations. Indeed, substituting *a*_11_ = *a*_22_ = *a, a*_12_ = *a* − *ξ* and *a*_21_ = *a* + *κ* in (5, 7), we obtain the conditions for the stable coexistence of both populations with unbalanced gene transfer in the oscillating environment:

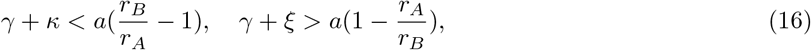

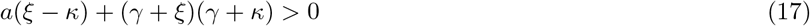

The conditions (16) and (17) are the counterparts of (12) and (13), respectively, for the case of unbalanced gene transfer in a fixed environment. From (16,17) and (8), together, it follows that the environmental variations can change the composition and the total abundance of the two populations in the coexisting equilibrium. Environmental oscillations destroy the coexistence of the two populations observed in the fixed environment if any of the conditions (16,17) is violated, but (8) holds.

The limitations of the model are discussed in the Supplementary Information. There, we compare the solutions obtained by numerically integrating the system (3,4) with time-dependent rates and the coarse grained counterparts obtained via (9,10). The comparison is based on the Euclidean distance of the time-averaged fractions of both populations obtained via (3,4) and those obtained from (9,10). The difference is negligible for the regions IV, V, VI, but not for the coexistence regimes. Nevertheless, even in the worst case scenario, both methods show the coexistence of both populations although the fractions of the two populations at equilibrium differ. We also compare the relative error of total population abundances. The relative error increases as one approaches the boundary *a*(*ξ* −*κ*) + *ξκ* = 0 where 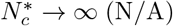, thus increasing the discrepancy between the results of the coarse-grained approach and the time average of the solution obtained via time-dependent rates, which remains bounded in general.

## DISCUSSION

In this work, we developed a mathematical model to explore the role of HGT in the interaction between two cohabiting populations in an oscillating vs a fixed environment. We explored what seems to be the simplest possible model in which two populations differed by a single gene allele, such that each entity in the pool carried only one allele of the given gene that is subject to HGT through copying the gene in the donor cell, and then, substituting the existing gene of the recipient cell, which corresponds to homologous recombination [14, 18]. Notwithstanding the ultimate simplicity of this scheme, it is biologically realistic, for example, in the context of acquisition of multi-drug-resistance [64]. We start with the stochastic description of all possible interactions between two populations, namely, reproduction and death of each population, competition within and between populations, and gene transfer between the populations. Then, we take the continuous limit and obtain the deterministic description of the time variations of the size of each population (see Additional File1, [53, 54]). The dynamical system obtained by this procedure is a variation of the well-known replicator dynamics [55–59], with varying total abundance of both populations. The composition and the total abundance of both populations in the equilibrium states are found from the rest points of this system.

By examining this mathematical model, we show that stable coexistence of the two populations in a fixed environment is unattainable within the assumption that the gene allele replacement via HGT only affects the reproduction rate, the competing abilities of both populations are the same within and between populations, and HGT between the populations is balanced, that is, there is no preferred direction in the gene flow between the populations. These results reflect the classic competitive exclusion (Gauze) principle according to which, in a spatially homogeneous environment, a population with an inferior reproduction rate will inevitably go extinct when competing with a fitter population for the same resource, within the same niche [60–62]. Notable, however, if the HGT is unbalanced, that is, the rates of gene transfer in the two directions are unequal, the exclusion principle does not apply anymore, and the two populations can coexist.

Evolutionary outcomes in a time-varying environment can drastically differ from those in the fixed environment, and despite the simplicity of our model, the emerging dynamics is complex, with the outcomes critically depending on the parameter combination. Environmental oscillations are incorporated into the model by assuming that the reproduction and gene transfer rates are time-periodic functions, with the implicit assumption that these rates are affected by the changes in the environment. The oscillations were set up such that the averages across environmental variations coincided with the constant rates in the fixed environment, providing for a fair comparison of the dynamics in the two types of environment.

The environmental oscillations occur on a fast time scale, whereby the populations are exposed to numerous changes in the environment before approaching any potential equilibrium. We adopted a coarse-graining approach to account for the environmental variations whereby the solution for the composition and total abundance of both populations was obtained as a combination of two components, one delineating the slow-time (coarse grained) behavior and the other capturing the oscillating component [44, 45]. By averaging over the period of environmental variations and retaining the first two leading-order terms of the varying quantities, we obtained the replicator dynamics model with varying total abundance that features fitness contributions derived from two distinct games. The first game represents the fixed/averaged environment whereas the second one arises due to environmental fluctuations, with the payoffs in this game determined by the intersection of the oscillating components of the reproduction and gene transfer equilibrium rates. These new payoffs are such that the fitness of a particular population is defined by the average of the product of the competitor’s reproduction rate and the gene transfer equilibrium rate between the populations throughout environmental oscillations. For the emergence of the new game, reproduction and gene transfer balance rates must be differentially affected by the environment. Notably, variations either in the reproduction rates or in the gene transfer rate alone did not result in new evolutionary scenarios compared to the fixed environment case.

The emerged game maintains a straightforward payoff structure: once the environmental variations of reproduction and gene transfer rates are given, then either one of the strategies consistently dominates, or there exists a unique non-trivial equilibrium for any given total abundance of both populations. With these adjusted fitness terms, stable coexistence between the two populations becomes possible within a broad range of model parameters. Moreover, depending on the combination of the time-dependent reproduction and gene transfer balance rates, the co-existence of the two populations manifests as all major types of symbiotic relationships. Two populations can coexist in a purely competitive symbiosis, where the fitness of each is negatively impacted by the presence of the other population; a host-parasite relationship, whereby the impact of the second population is positive for one and negative for the other population; and in mutualistic symbiosis, where the presence of the other population is reciprocally beneficial.

We further analyzed the behavior of the total abundance of both populations and defined the regions of synergistic interactions between the competing populations. In this case, synergy is observed, that is, the total abundance of the two populations at the stable coexistence in the oscillating environment is greater than the total abundance in the fixed environment, that is, the population size of the winner at equilibrium, under the same model parameter values. As could be expected, mutualism necessarily entails synergy and yields the greatest total abundance of both populations among all the regimes by increasing the resource utilization in the environment. Notably, however, under certain combinations of the reproduction and HGT rates, the synergistic effect was observed also in the cases of parasitism and even pure competitive symbiosis.

Outside of the coexistence regimes, the outcomes of the competition between two populations in a stable environment can persist in the oscillating environment such that the winner in the fixed environment wins in the oscillating environment as well. However, in another region of the phase space, the outcome can be reversed due to environmental oscillations. Moreover, there is also a bistability regime where one or the other population goes extinct depending on the initial conditions, a scenario precluded in the fixed environment assuming equal competing abilities and balanced gene transfer. Despite our prior discussion on coexistence and potential outcomes in an oscillating environment with balanced gene transfer in the fixed environment, the introduction of oscillations can notably alter the composition at the coexistence equilibrium, even if both populations coexist in the fixed environment due to non-zero gene transfer balance. Environmental variations, in this case, can even destroy the stable coexistence of the two populations, attained in the fixed environment with unbalanced HGT.

From the methods point of view, it is worth pointing out that coarse graining was an essential ingredient of the present analysis because without it, the symbiotic relationships between two populations would be impossible to elucidate because the fitness gradients can change their signs throughout environmental oscillation due to the oscillating rates.

From the biological standpoint, environmental oscillations provide for the synergy between the two populations by lowering the intensity of the inter-population competition below the level of the intra-population competition. This work shows that stabilization of strain diversity and increase of ecological complexity via HGT in an oscillating environment is an intrinsic feature of even the simplest microbiomes that emerges under minimal assumptions on the basic processes occurring within a microbial community. Given that both environmental variation and HGT are ubiquitous phenomena that affect any microbiome [65, 66], these conclusions appear to be broadly applicable. Notably, all emerging coexistence regimes provide for synergy between the populations indicating that HGT in an oscillating environment is favorable for a microbial community perceived as an integral whole.

Evidently, the model used in this work is grossly (and deliberately) over-simplified. Real microbiomes encompass interactions among thousands of microbial strains and species, HGT of multiple genes via different routes and many other processes [14, 50, 66]. Nevertheless, the general principle established here should apply, amplified by the microbiome complexity. Characterization of the conditions for stabilization of microbiome diversity and the factors that can perturb it is crucial for understanding the role of the microbiome in health and diseases as well as the ecology of microbial communities.

## SUPLLEMENTARY INFORMATION

The manuscript contains supplementary material. Additional File1 contains derivations and analysis of (1,2) and (9,10), and limitations of the proposed model.

## ACKNOWLEDGEMENTS

The authors are thankful to Armen E. Allahverdyan and Koonin group members for valuable discussions.

## AUTHORS’ CONTRIBUTIONS

SGB initiated the project; SGB, YIW and EVK designed the model; SGB implemented the model; SGB, SKG, YIW and EVK analyzed the results; SGB and EVK wrote the manuscript that was read, edited and approved by all authors.

## FUNDING

The authors are supported through the Intramural Research Program of the National Library of Medicine, National Institutes of Health.

## AVAILABILITY OF DATA AND MATERIALS

This is a theoretical work that did not involve data analysis.

## ETHICS APPROVAL AND CONSENT OF PARTICIPATE

Not applicable.

## COMPETING INTERESTS

The authors declare they have no competing interests.

## Notes

### Competing Interest Statement

The authors have declared no competing interest.

