## Supplementary Material for "Microbial diversity and ecological complexity emerging from environmental variation and horizontal gene transfer in a simple mathematical model"

### I. COMPETITION IN THE FIXED ENVIRONMENT.

#### A. Derivation of the time-behavior of two population sizes from elementary processes.

In this section we are going to derive the deterministic equations governing the populations sizes $n(t)$ and $m(t)$ of $A$ and $B$ populations, respectively, from the more elementary processes. The derivation is based on the approach developed in [1, 2] We assume that the carrying capacity of the environment is fixed $K = const$. The available resources in the environment at time $t$ is $K - n(t) - m(t)$. We assume that in each small time interval $\Delta t$, either one or two entities are selected. These selections are governed by probabilities $\mu$ and $1-\mu$, respectively. Specifically, for type $A$ (and analogously for $B$), the elementary processes are as follows:

$$\begin{aligned}
&\text{Processes with two entities} \\
&AE \to AA, \quad \text{reproduction,} \\
&AA \to AE, \quad \text{intraspecies competition,} \\
&AB \to EB, \quad \text{interspecies competition,} \\
&AB \to AA, \quad \text{gene trasfer A} \to \text{B,} \\
&\text{One entity process} \\
&A \to \emptyset, \quad \text{death of A}
\end{aligned} \tag{1}$$

We denote the probability of encountering $n(t)$ units of type $A$ and $m(t)$ units of type $B$ at time $t$ as $P(n, m, t)$. The temporal evolution of $P(n, m, t)$ follows the Markov dynamics, characterized by the following transition rates:

$$\begin{aligned}
T(n+1, m|n, m) &= 2\mu\rho_A \frac{n}{K}\frac{K-n-m}{K-1}, \\
T(n-1, m|n, m) &= \mu c_{AA} \frac{n}{K}\frac{n-1}{K-1} + 2\mu c_{AB} \frac{n}{K}\frac{m}{K-1} + (1-\mu)\delta_A \frac{n}{K}, \\
T(n, m+1|n, m) &= 2\mu\rho_B \frac{m}{K}\frac{K-n-m}{K-1}, \\
T(n, m-1|n, m) &= \mu c_{BB} \frac{m}{K}\frac{m-1}{K-1} + 2\mu c_{BA} \frac{m}{K}\frac{n}{K-1} + (1-\mu)\delta_B \frac{m}{K}, \\
T(n+1, m-1|n, m) &= 2\mu\gamma_{AB} \frac{n}{K}\frac{m}{K-1}, \\
T(n-1, m+1|n, m) &= 2\mu\gamma_{BA} \frac{m}{K}\frac{n}{K-1},
\end{aligned} \tag{2}$$

Here $\rho_A$ and $\delta_A$ are reproduction and death rates respectively. $c_{AA}$, $c_{AB}$, and standing for intra and inter-species competition, $\gamma_{AB}$ is the gene transfer rate from $B \to A$ once $A$ and $B$ are chosen. to get the intuition, let us consider the transition rate $T(n+1, m|n, m)$, that is the reproduction of $A$. The rate is defined as follows, with probability $\mu$ two ball process is chosen, then with probability $n/K$ an entity of $A$ type and $K - n - m/K - 1$ an entity of $E$ is chosen, and finally with $\rho_A$ the birth happens.

The master equation for $P(n, m, t)$ has the following form

$$\frac{d}{dt}P(n, m, t) = \sum_{s'} T(s|s')P(s', t) - T(s'|s)P(s, t), \tag{3}$$
$$\text{here} \quad s = \{n, m\}\,, \; S' = \{n-1, m+1\}\,, .., \{n+1, m-1\}\,.$$

We multiply (3) by $n$ and average over possible values of $n, m \in [0, K]$. That is defining the average $< n >= \sum_{n,m=0}^{K} nP(n, m, t)$ we get from (3)

$$\frac{d}{dt} < n > \; = \sum_{n,m=0}^{K} \sum_{S''} n(T(s|s'')P(s'', t) - T(s''|s)P(s'', t)) + \sum_{n,m=0,K} \sum_{S'/S''} n(T(s|s')P(s', t) - T(s, |s)P(s', t)) \tag{4}$$

here $S'' = \{\{n, m+1\}\,, \{n, m-1\}\}$ is the set of states that include variations in $m$ only, the sum over these states nullifies in. Indeed, from (4) we get

$$\sum_{n,m=0}^{K}\sum_{S''} n(T(s|s'')P(s'',t)-T(s''|s)P(s'',t)) = \sum_{n,m=0}^{K} nT(n,m|n,m+1)P(n,m+1) - \tag{5}$$
$$\sum_{n,m=0}^{K} nT(n,m+1|n,m)P(n,m) + \sum_{n,m=0}^{K} nT(n,m|n,m-1)P(n,m-1) - \sum_{n,m=0}^{K} nT(n,m-1|n,m)P(n,m)$$

Changing the variable in the summation $m = m' - 1$ in the first term and $m = m' + 1$ in the third one, and noting that $T(n,-1|n,0) = T(n,0|n,-1) = T(n,K|n,K+1) = T(n,K+1|n,K) = 0$ we get that the right hand-side nullifies.

For the states in $S'/S''$ we do the same trick, that is changing the summation variables for $n$ and $m$. We further assume that the averages in the form $< nm >=< n >< m >$, that is the mean-field approach, and finally neglecting the terms of $1/K$ in intra and inter species transition rates, that is $\frac{n}{K}\frac{n-1}{K-1} = \frac{n^2}{K(K-1)}$ (repeating the same procedure for $B$), we get the deterministic equations governing the population sizes $n(t) =< n(t) >$ and $m(t) =< m(t) >$

$$\frac{dn}{dt} = n(r_A - a_{11}n - a_{12}m) + \gamma mn, \tag{6}$$
$$\frac{dm}{dt} = m(r_B - a_{21}n - a_{22}m) - \gamma mn, \tag{7}$$

where $r_A =< \mu\frac{2\rho_A}{K-1} - (1-\mu)\frac{\delta_A}{K} >$, $a_{11} =< \mu\frac{2\rho_A + c_{AA}}{K(K-1)} >$, $a_{12} =< \frac{2\mu c_{AB}}{K(K-1)} >$ and $\gamma = 2\mu\frac{\gamma_{AB}-\gamma_{BA}}{K(K-1)}$, here $<>$ are standing for averaging over $P(n,m)$. Those for $B$ are obtained in similar way.

From (6,7) one can derive the equivalent representation of the evolutionary processes in terms of the total abundance of both populations $N(t) = n(t) + m(t)$ and the fractions of $A$ and $B$ populations in the total abundance $p_A = \frac{n(t)}{N(t)}$ and $p_B = \frac{m(t)}{N(t)}$, represented in the (3,4) in the main text.

#### B. Dynamical analysis of (6,7)

The two representations (6,7) and in terms of $p_A(t)$ and $N(t)$ are equivalent, however, the analysis is more easily accomplished in the system (6,7) due to the non-dimensialization of the system [3]. The non-dimensialzation of the system is done as follows

$$\frac{du}{dt_1} = u(1 - u - \alpha v), \tag{8}$$
$$\frac{dv}{dt_1} = \beta v(1 - \delta u - v), \tag{9}$$

where $t_1 = r_A t$ $u = \frac{a_{11}}{r_A}n$, $v = \frac{a_{22}}{r_B}m$, $\beta = \frac{r_B}{r_A}$, $\alpha = \frac{a_{12}-\gamma}{r_A}\frac{r_B}{a_{22}}$ and $\delta = \frac{a_{21}+\gamma}{r_B}\frac{r_A}{a_{11}}$.

The possible equilibrium states of (8,9) are $(u^*, v^*) = (0,0)$, $(1,0)$, $(0,1)$, $(\frac{1-\alpha}{1-\alpha\delta}, \frac{1-\delta}{1-\alpha\delta})$.

Note, that $\text{sgn}(1-\alpha\delta) = \text{sgn}(\text{Det}\mathcal{A})$, where $\mathcal{A}$ is the following community matrix

$$\mathcal{A} = \begin{bmatrix} a_{11} & a_{12} - \gamma \\ a_{21} + \gamma & a_{22} \end{bmatrix} \tag{10}$$

The equilibrium state is stable if the eigenvalues of the Jacobian matrix of the dynamical system in the equilibrium have negative real parts $\text{Re}\lambda_i < 0,\ i = 1,2$ [3, 4]. The Jacobian matrix of the dynamical system (8,9) has the following form

$$J|_{u^*,v^*} = \begin{bmatrix} 1 - 2u^* - \alpha v^* & -\alpha u^* \\ -\beta\delta v^* & \beta(1 - 2v^* - \delta u^*) \end{bmatrix} \tag{11}$$

The eigenvalues have negative real parts if $\text{Tr}J|_{u^*,v^*} < 0$ and $\text{Det}J|_{u^*,v^*} > 0$.

For $(u^*, v^*) = (0,0)$ we get both $\mathrm{Tr}J|_{0,0} > 0$ and $\mathrm{Det}J|_{0,0} > 0$, hence the equilbrium is unstable, as long as $\beta > 0$ which is by default assumption always holds.

In the equilibrium $(u^*, v^*) = (1,0)$ we obtain $\mathrm{Tr}J|_{1,0} = -1 + \beta(1-\delta)$ and $\mathrm{Det}J|_{1,0} = -\beta(1-\delta)$. Thus, the equilibrium is stable if $\delta > 1$. Similarly, we obtain the following condition $\alpha > 1$ for the stability of $(u^*, v^*) = (0,1)$.

The nontrivial equilibrium $(u^*, v^*) = (\frac{1-\alpha}{1-\alpha\delta}, \frac{1-\delta}{1-\alpha\delta})$ is meaningful if $u^* > 0,\ v^* > 0$, since $n = r_A/a_{11}u^*$ and $m^* = r_B/a_{22}v^*$. The condition $(u^*, v^*) > 0$ holds if either $\alpha < 1,\ \delta < 1,\ \alpha\delta < 1$ or $\alpha > 1,\ \delta > 1,\ \alpha\delta > 1$. For the stability of the equilibrium we get

$$\mathrm{Tr}J|_{u^*,v^*} = -(u^* + \beta v^*), \quad \mathrm{Det}J|_{u^*,v^*} = \beta u^* v^* (1-\alpha\delta) \tag{12}$$

$\mathrm{Tr}J|_{u^*,v^*} < 0$ as long as $u^* > 0,\ v^* > 0$ that is for both $\alpha < 1,\ \delta < 1,\ \alpha\delta < 1$ and $\alpha > 1,\ \delta > 1,\ \alpha\delta > 1$ cases. However, the $\mathrm{Det}J|_{u^*,v^*} > 0$ only for the first case, that is $\alpha < 1,\ \delta < 1,\ \alpha\delta < 1$. Thus, the non-trivial equilibrium is either stable–representing coexistence of both population, or unstable–separating the basins of attraction of $(u^*, v^*) = (1,0)$ and $(u^*, v^*) = (0,1)$, if exists. Note that, in the latter case both $(u^*, v^*) = (1,0)$ and $(u^*, v^*) = (0,1)$ are still stable, while, in the former case they are unstable.

In the main text, we focus on the situation where all competition rates are equal $a_{ij} = a$ and $\gamma = 0$. In this case we get $\alpha = \frac{r_B}{r_A}$ and $\delta = \frac{r_A}{r_B}$, thus, if $r_A \neq r_B$, then there is no coexistence of both populations, even more there is no bistability either. The only stable equilibrium state in this case is either $(1,0)$ or $(0,1)$, depending on the reproduction rates $r_A$ and $r_B$. The, one with greater reproduction rate outcompetes the opponent.

In the case $\alpha < 1,\ \delta < 1,\ \alpha\delta > 1$, where $u^* < 0, v^* < 0$ the solution of (8,9) go infinity. Indeed, let us take the particular case $\alpha = \delta$ and $\beta = 1$, and let's assume that $u(0) = v(0)$. Then, we get from (8,9) the following equation

$$\frac{du}{dt} = u(1 - (1+\alpha)u) \tag{13}$$

Now, we get $\alpha < 1$ and $\alpha^2 > 1$, which is possible if $\alpha < -1$, thus, $1 + \alpha < 0$. Denoting $\zeta = |1+\alpha| > 0$, and solving for $u$, by assuming that $1/u(t_f) \to 0$, that is assuming that the solution goes to infinity, one can find the time of this divergence $t_f = \ln \frac{1+\zeta u_0}{\zeta u_0}$, that is finite for all $u_0 > 0$.

### II. OSCILLATIONS IN THE ENVIRONMENT: COARSE-GRAINED DESCRIPTION

In the fixed environment the fractions and total abundance of both populations are found by the following system of differential equations

$$\frac{dp_i}{dt} = p_i(f_i - \sum_j p_j f_j), \quad i,j = A,B \tag{14}$$

$$\frac{dN}{dt} = N \sum_j p_j f_j. \tag{15}$$

where $p_A + p_B = 1$. The fitnesses of the populations can be interpreted as expected payoffs in the following game

$$\begin{bmatrix} r_A - a_{11}N & r_A - a_{12}N + \gamma N \\ r_B - a_{21}N - \gamma N & r_B - a_{22}N \end{bmatrix} \tag{16}$$

Thus, the fitness of $A$ and $B$ populations are equal to $f_A = p_A(r_A - a_{11}N) + (1-p_A)(r_A - a_{12}N + \gamma N)$ and $f_B = p_A(r_B - a_{21}N - \gamma N) + (1-p_A)(r_B - a_{22}N)$, where we have taken into account $p_A + p_B = 1$. Note, that the total abundance of both populations proliferates with by the mean fitness of both populations. In the equilibrium, the fitnesses of each population, presented in the equilibrium, nullifies.

We now proceed to the derivation of coarse-grained behavior of the fractions and total abundance of both populations in oscillating environment.Though, it has to be noted that the same results could be obtained by considering the population sizes $n(\tau), m(\tau)$ described by (6,7).

We assume the environment is periodically varying with frequency $\omega > 1$ and period $2\pi$. We assume the environment oscillates in the fast time scale $\tau = \omega t$.

The variation in the environment induce oscillations in the reproduction rates and gene transfer balance rate, that is

$$r_A(\tau+2\pi) = r_A(\tau), \quad r_B(\tau+2\pi) = r_B(\tau), \quad \gamma(\tau+2\pi) = \gamma(\tau), \tag{17}$$

while the within and between-competition rates are time independent $a_{ij} = \text{const}$.

We will further assume that the time dependent rates can be represented as a sum of constant and oscillating parts $r_i(\tau) = r_i + \tilde{r}_i$, $i = A, B$, such that $\overline{r_i(\tau)} \equiv \int_0^{2\pi} \frac{d\tau}{2\pi} r_i(\tau) = r_i$, and $\overline{\gamma(\tau)} = \gamma$.

Thus, in terms of the average rates we recover the deterministic description of the model in the constant environment (14,15) in the main text.

The fitnesses of each population in the oscillating environment can be represented as sum of terms with the mean rates and oscillating parts, for population $A$ these are

$$f_A(p(\tau), N(\tau)) = p(\tau)(r_A - a_{11}N(\tau)) + (1-p(\tau))(r_A - a_{12}N(\tau) + \gamma N(\tau)), \tag{18}$$
$$\tilde{f}_A(p(\tau), N(\tau)) = \tilde{r}_A + (1-p(\tau))\tilde{\gamma}N(\tau) \tag{19}$$

here we used the normalizations $p_A(\tau) + p_B(\tau) = 1$. We will discuss only for $p_A(\tau) = p(\tau)$. The case for $B$ can be easily recovered then.

The representations of the fitnesses as a sum of two terms allow us to rewrite the right hand-side of (14) as a sum of two term $G(p(\tau), N(\tau)) + \tilde{G}(p(\tau), N(\tau))$, where the last term includes only oscillating rates. Similarly, for (15) we represent the right hand-side as a $F(p(\tau), N(\tau)) + \tilde{F}(p(\tau), N(\tau))$.

We seek the solution of (14,15) with time-dependent rates as a sum of slowly-varying (varying in $t$) and oscillating parts (varying in $\tau$)

$$p = p(t) + \pi(p(t), N(t), \tau), \tag{20}$$
$$N = N(t) + \mathcal{V}(p(t), N(t), \tau) \tag{21}$$

The oscillating parts averages to zero $\overline{\pi(p(t), N(t), \tau)} = \overline{\mathcal{V}(p(t), N(t), \tau)} = 0$. Now we put (20,21) into the equations, and expand the right-hand side over $\pi(p(t), N(t), \tau)$ and $\mathcal{V}(p(t), N(t), \tau)$

$$\dot{p}(t) + \dot{p}\frac{\partial}{\partial p}\pi + \dot{N}\frac{\partial}{\partial N}\pi + \omega\partial_\tau\pi = G(p,N) + \tilde{G}(p,N) + \pi\frac{\partial}{\partial p}(G+\tilde{G}) + \mathcal{V}\frac{\partial}{\partial N}(G+\tilde{G}), \tag{22}$$
$$\dot{N} + \dot{p}\frac{\partial}{\partial p}\mathcal{V} + \dot{N}\frac{\partial}{\partial N}\mathcal{V} + \omega\partial_\tau\mathcal{V} = F(p,N) + \tilde{F}(p,N) + \pi\frac{\partial}{\partial p}(F+\tilde{F}) + \mathcal{V}\frac{\partial}{\partial N}(F+\tilde{F}). \tag{23}$$

The oscillating parts are searched for via expanding by $\frac{1}{\omega}$, that is $\pi = \sum_{k=1}^{\infty} \frac{1}{\omega^k}\pi_k$ and $\mathcal{V} = \sum_{k=1}^{\infty} \frac{1}{\omega^k}\mathcal{V}_k$. Note, that the averages over the period of environmental variations is again zero for $\pi_k, \mathcal{V}_k$. We insert them back into (22,23), and keep the terms up to $\frac{1}{\omega^2}$. The obtained equations containt terms in various order of magnitude and varying on different time scales. We first collect the terms of $O(1)$ that vary on the fast time. These are

$$\partial_\tau\pi_1 \ = \ \tilde{G} \equiv p\big(\tilde{r}_A + \tilde{\gamma}(1-p)N - (p\tilde{r}_A + (1-p)\tilde{r}_B)\big) \tag{24}$$
$$\partial_\tau\mathcal{V}_1 \ = \ \tilde{F} \equiv N(p\tilde{r}_A + (1-p)\tilde{r}_B). \tag{25}$$

These equations could be directly integrated, since the left hand side depend of both equations depend on $\tau$ through the oscillating rates. Denoting the primitives of reproduction and gene transfer balance rates as $\hat{r}_i$, $i = 1,2$ and $\hat{\gamma}$, that is $\partial_\tau\hat{r}_i = \tilde{r}_i$ and $\partial_\tau\hat{\gamma} = \tilde{\gamma}$, then we get $\pi_1 = \hat{G}$ and $\mathcal{V}_1 = \hat{F}$, where $\hat{G}$ and $\hat{F}$ are the right hand sides of (24,25) with primitives of the rates. Now, we average (22,23) over the period of oscillations, note that the terms $\tilde{G}$ and $\tilde{F}$. Also, the following terms nullify $\pi_1\frac{\partial}{\partial p}G$, $\pi_1\frac{\partial}{\partial p}F$, $\mathcal{V}_1\frac{\partial}{\partial N}G$ and $\mathcal{V}_1\frac{\partial}{\partial N}F$, because only $\pi_1$, $\tau$ depend on $\tau$, and these have zero averages. Similarly, on the left hand side the terms of the form $\dot{p}\frac{\partial}{\partial_p}\pi$, also nullify after averaging.

Thus, we end up with the following set of differential equations

$$\frac{dp}{dt} \ = \ G(p,N) + \frac{1}{\omega}\overline{\pi_1\frac{\partial}{\partial p}\tilde{G}} + \frac{1}{\omega}\overline{\mathcal{V}_1\frac{\partial}{\partial N}\tilde{G}}, \tag{26}$$
$$\frac{dN}{dt} \ = \ F(p,N) + \frac{1}{\omega}\overline{\pi_1\frac{\partial}{\partial p}\tilde{F}} + \frac{1}{\omega}\overline{\mathcal{V}_1\frac{\partial}{\partial N}\tilde{F}} \tag{27}$$

In the averaging process, we use the solutions of (24,25). Note, the following identities between the oscillating rates $\overline{\hat{h}_1\tilde{h}_1} = 0$ and $\overline{\hat{h}_1\tilde{h}_2} = -\overline{\tilde{h}_1\hat{h}_2}$, where $h_1, h_2$ stand for reproduction and gene transfer balance rates. Using the last observation, one can show that the diagonal terms $\overline{\pi_1\frac{\partial}{\partial p}\tilde{G}} = \overline{\mathcal{V}_1\frac{\partial}{\partial N}\tilde{F}} = 0$. Let us calculate the non-zero terms

$$\begin{aligned}\overline{\pi_1\frac{\partial}{\partial p}\tilde{F}} = \overline{\hat{G}\frac{\partial}{\partial p}\tilde{F}} &= \overline{p\big(\hat{r}_A + \hat{\gamma}(1-p)N - p\hat{r}_A - (1-p)\hat{r}_B\big)N(\tilde{r}_A - \tilde{r}_B)} = \\ &= \overline{Np\big(\hat{r}_A(\tilde{r}_A - \tilde{r}_B) + \hat{\gamma}(\tilde{r}_A - \tilde{r}_B)N(1-p) - p\hat{r}_A(\tilde{r}_A - \tilde{r}_B) - (1-p)\hat{r}_B(\tilde{r}_A - \tilde{r}_B)\big)} = \\ &= \overline{Np(1-p)\big((\hat{r}_A - \hat{r}_B)(\tilde{r}_A - \tilde{r}_B) + \hat{\gamma}(\tilde{r}_A - \tilde{r}_B)N\big)} = N^2p(1-p)(\overline{\hat{r}_B\tilde{\gamma}} - \overline{\hat{r}_A\tilde{\gamma}}).\end{aligned} \tag{28}$$

here we have used the above identities. The remaining non-zero term is found in similar way

$$\overline{\mathcal{V}_1\frac{\partial}{\partial N}\tilde{G}} = \overline{\hat{F}\frac{\partial}{\partial N}\tilde{G}} = \overline{N(p\hat{r}_A + (1-p)\hat{r}_B)\tilde{\gamma}p(1-p)} = Np(1-p)(p\overline{\hat{r}_A\tilde{\gamma}} + (1-p)\overline{\hat{r}_B\tilde{\gamma}}). \tag{29}$$

Putting these terms back into the system (26,27) we obtain

$$\frac{dp_A}{dt} = p_A(f_A - \sum_{j=A,B} p_j f_j) + Np(1-p)(p\kappa + (1-p)\xi), \tag{30}$$

$$\frac{dN}{dt} = N\sum_{j=A,B} p_j f_j + N^2 p(1-p)(\xi - \kappa). \tag{31}$$

where we have denoted $\xi = \frac{1}{\omega}\overline{\hat{r}_B\tilde{\gamma}}$ and $\kappa = \frac{1}{\omega}\overline{\hat{r}_A\gamma}$. Note, that the fitnesses $f_A, f_B$ in (30,31) are the same as in the fixed/averaged environment.

The added terms could be represented in the form of additional fitnesses. That is

$$p(1-p)(\phi_A - \phi_B) = Np(1-p)(p\kappa + (1-p)\xi), \tag{32}$$

$$p\phi_A + (1-p)\phi_B = Np(1-p)(\xi - \kappa). \tag{33}$$

Solving these equations for $\phi_A, \phi_B$ we find that $\phi_A = N\xi(1-p)$ and $\phi_B = -N\kappa p$.

Putting these back into (30,31) we derive

$$\frac{dp_i}{dt} = p_i(f_i + \phi_i - \sum_j p_j(f_j + \phi_j)), \quad i,j = A,B \tag{34}$$

$$\frac{dN}{dt} = N\sum_j p_j(f_j + \phi_j). \tag{35}$$

The initial conditions for (34,35) are found from (20,21). That is we find the initial condition $p_0, N_0$ by solving the following system of equations with respect to $p_0, N_0$,

$$p(0) = p_0 + \frac{1}{\omega}\hat{G}(0), \tag{36}$$

$$N(0) = N_0 + \frac{1}{\omega}\hat{F}(0) \tag{37}$$

with obvious constraints $0 \le p_0 \le 1, N_0 > 0$, here $p(0), N(0)$ are the initial condition for $p(\tau), N(\tau)$.

Note, that the (34,35) could be obtained from (14,15) by substituting $a_{12} \to a_{12} - \xi$ and $a_{21} \to a_{21} + \kappa$. This holds for (6,7), also. That is, if we derive the coarse-grained behavior through $n, m$ we end up in (6,7), with $a_{12} \to a_{12} - \xi$ and $a_{21} \to a_{21} + \kappa$.

Thus, dynamical analysis of the (IB) holds with these substitution.

For example, we obtained that coexistence is possible if $\alpha < 1$,$\delta < 1$ and $\alpha\delta < 1$. Now we substitute $a_{12}$ and $a_{21}$ by $a_{12} - \xi$ and $a_{21} + \kappa$, respectively, and end up with the following conditions

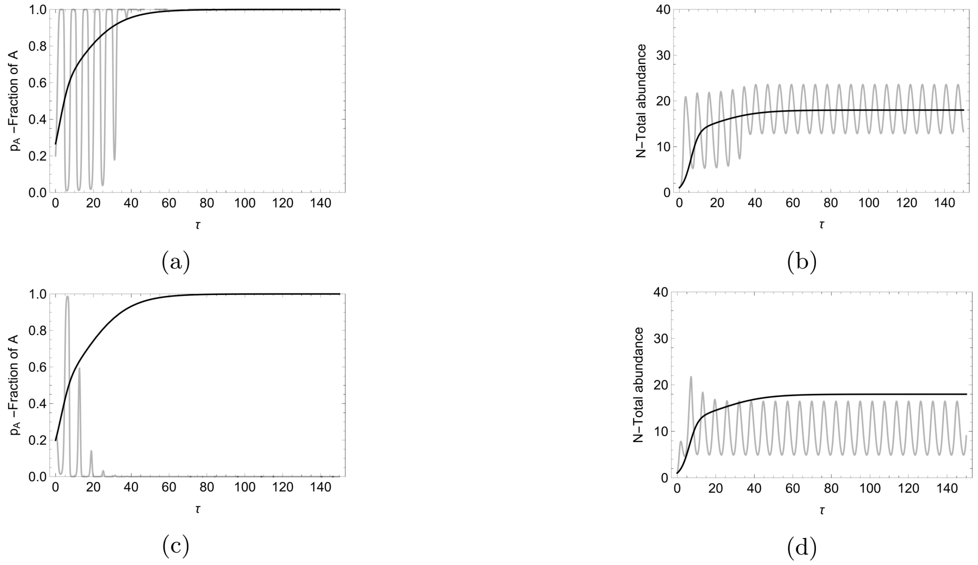


FIG. 1: **Comparing the behavior of the fractions and total abundance of both populations for $\gamma$ and $-\gamma$.** In the first and last row $\gamma = \frac{1}{2}\sin\tau$ and $\gamma = -\frac{1}{2}\sin\tau$. The reproduction rates are as follows $\tilde{r}_A = -0.36\cos\tau + 0.5\sin\tau$, $\tilde{r}_B = -0.84\cos\tau$ and $\tilde{r}_A = 0.36\cos\tau + 0.5\sin\tau$, $\tilde{r}_B = 0.84\cos\tau$ for the top and bottom rows, respectively. For both cases $\xi = -0.07$ and $\kappa = -0.03$. The remaining model parameters are $r_A = 1.8$, $r_B = 1$, $a = 0.1$ and $\omega = 3$.

$$\frac{a_{12} - \xi - \gamma}{a_{22}}\frac{r_B}{r_A} < 1, \quad \frac{a_{21} + \gamma + \kappa}{a_{11}}\frac{r_A}{r_B} < 1, \qquad (38)$$
$$\frac{a_{12} - \xi - \gamma}{a_{22}}\frac{a_{21} + \gamma + \kappa}{a_{11}} < 1.$$

Particularly, for the $\gamma = 0$ and $a_{ij} = a$ case we get the conditions discussed in the main text

$$(1 - \frac{\xi}{a}) < \frac{r_A}{r_B}, \quad (1 + \frac{\kappa}{a}) < \frac{r_B}{r_A}, \qquad (39)$$

$$(1 - \frac{\xi}{a})(1 + \frac{\kappa}{a}) < 1 \qquad (40)$$

### III. NUMERICAL VALIDATION AND LIMITATION OF THE MODEL.

As we have discussed in the Section I B, the coarse grained approach failed in the regime where (39) is valid, but (40) holds in reverse. In this case, the community matrix $\mathcal{A}$, representing the competition rates within and between populations has negative determinant. Indeed, the community matrix has the following form

$$\mathcal{A} = \begin{bmatrix} a & a - \xi \\ a + \kappa & a \end{bmatrix} \qquad (41)$$

from which one get that Det$\mathcal{A} = a(\xi - \kappa) + \xi\kappa$, then we get Det$\mathcal{A} < 0$ if (40) holds with reverse inequality sign. Note, that the DetA stands in the denominator of the total abundance of both populations in the coexisting regime $N_c^*$. Indeed, as one get closer to the $a(\xi - \kappa) + \xi\kappa = 0$ line as greater becomes $N_c^*$, thus increasing the discrepancy between the results of the coarse-grained approach and the time average of the solution obtained via time-dependent rates. Before going further into the details, let us focus on the other limitation of the model.

The new payoff terms are determined via the averages of the product of gene transfer balance rate $\tilde{\gamma}$ and the primitives of the oscillating reproduction rate $\hat{r}_i$, $i = $ A, B, as it is seen from the form of new payoff terms $\xi = \frac{1}{\omega}\overline{\hat{r}_B\tilde{\gamma}}$ and $\kappa = \frac{1}{\omega}\overline{\hat{r}_A\gamma}$.

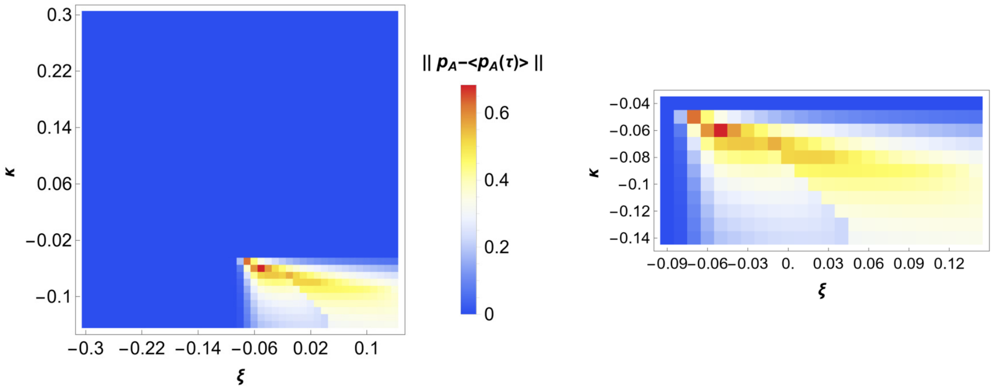


(a)

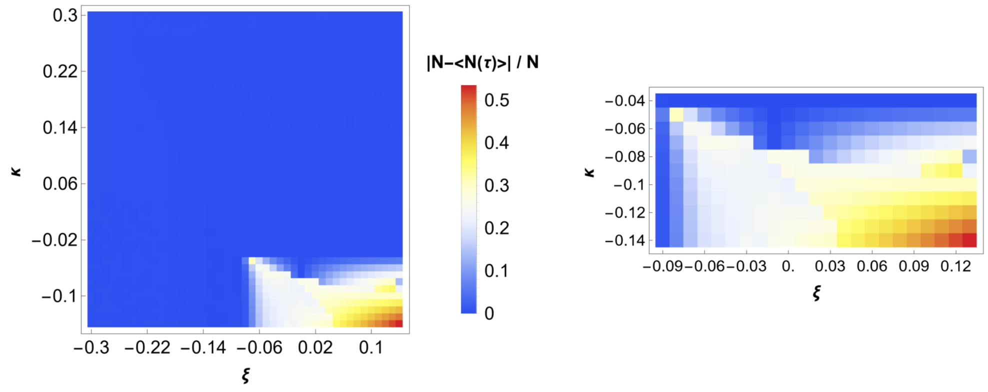


(b)

FIG. 2: **Discrepancies between the solutions obtained via coarse-grained approach (34,35) and (14,15) with time-dependent rates**. The top and bottom rows shows the discrepancy between the fractions and total abundance of both populations, respectively.Here $\gamma = z/2\sin\tau$, $\tilde{r}_A = a\cos\tau + 0.5\sin\tau$, $\tilde{r}_B = b\cos\tau$, where $a, b$ are found from the values of $\xi, \kappa$. The remaining model parameters are $T_f = 200$,$T_1 = 100$ $\omega = 3$, $a = 0.1$, $r_A = 1.8$ and $r_B = 1$.

Therefore, $\tilde{\gamma}$ and $-\tilde{\gamma}$ may provide the same payoffs $\xi$ and $\kappa$, as long as the sign of the reproduction rates are also changed. In terms of the coarse-grained approach these two cases provide the same outcome, however, the numerical solution of the model with time-dependent rates shows that the change in the sign of the rates may change the dynamics drastically, see Fig.(1). As it is seen from the figure, the coarse-grained behavior of the fraction and total abundance of both populations is in accordance with $\gamma = \frac{1}{2}\sin\tau$. Thus, the coarse-grained approach reveals the behavior of the oscillatory behavior of one of the possible signs, but not both.

To analyze further the limitation and discrepancy between the results obtained via coarse-grained approach and the solution of the model with time-dependent rates we introduce the following binary variable in the gene transfer balance $\tilde{\gamma} = zh(\tau)$, where $z = \pm 1$ and $h(\tau)$ is a periodic function.

Then for each $\xi$, $\kappa$ we calculate the following averages for both $z \pm 1$ and two different initial fractions $p' = 0.1, p'' = 0.9$, while $N(0) = \text{const}$. The corresponding initial conditions for coarse-grained approach are found from (36)

$$\langle p \rangle = \frac{1}{T_f - T_1}\int_{T_1}^{T_f} p(\tau)d\tau, \tag{42}$$

$$\langle N \rangle = \frac{1}{T_f - T_1}\int_{T_1}^{T_f} N(\tau)d\tau \tag{43}$$

where $p(\tau), N(\tau)$ are obtained via numerically solving (14,15). The limits of integration should be distinct enough to yield meaningful averaging. We compare these quantities with their counterparts obtained via coarse-grained approach at time $T_f$. As a measure of discrepancy we use Euclidean distance

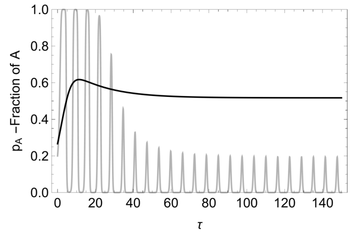


(a)

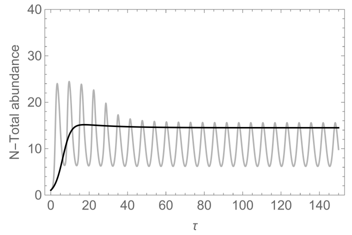


(b)

FIG. 3: **The fractions and total abundance of both populations corresponding to the maximum discrepancy in Fig.2.** Here $\xi = -0.05, \kappa = -0.06$, that are obtained for $\gamma = 1/2\sin\tau$ and $p(0) = 0.1$. Thus, $\tilde{r}_A = -0.72\cos\tau + 0.5\sin\tau$, $\tilde{r}_B = -0.6\cos\tau$ . The remaining model parameters are $\omega = 3$, $a = 0.1$, $r_A = 1.8$ and $r_B = 1$.

$$||p_A - \langle p_A\rangle|| = \sqrt{2(p_A - \langle p_A\rangle)^2},$$

and relative error $|N - \langle N\rangle|/N$ for the fractions and total abundance of both populations, respectively, here $p_A = p_A(T_f)$ and $N = N(T_f)$. Thus, the maximum error in the fractions is $\sqrt{2}$. We use relative error in the total abundance, since the average $\langle N\rangle$ remains bounded, in general, while $N(T_f)$ may be unbounded,as we have seen above.

The last step is we choose minimize the discrepancy between the fractions of both populations with respect to $z = \pm 1$ and $p(0) = (p', p'')$, that is

$$\min_{z=\pm 1, p', p''} ||p_A - \langle p_A\rangle|| \tag{44}$$

While, the relative errors of the total abundance of both populations are calculated at $z, p(0)$ for which the above minimization is obtained, for any given $\xi$ and $\kappa$.

The criterion (44) is used to construct the heat map of the euclidean distances presented in the Fig.2. As it is seen from the figure, the discrepancy between the coarse grained approach and the numerical solution obtained via time-dependent rates is negligible in the regions where either $A$ or $B$ wins. The discrepancy occurs in the coexistence regime, as one might expect. Comparing the relative error in the total abundance of both populations we see that it increases as one get closer to the line $a(\xi - \kappa) + \xi\kappa = 0$, as we have expected.

The worst case is presented in Fig.3, that is $\max_{\xi,\kappa}\min_{z\pm 1,p',p''}||p_A - \langle p_A\rangle||$. Even in the worst case, both methods represent coexistence between two populations, although the fractions are different.

---


[1] A. J. McKane and T. J. Newman, Physical review letters **94**, 218102 (2005).
[2] A. J. McKane and T. J. Newman, Physical Review E **70**, 041902 (2004).
[3] J. D. Murray, *Mathematical biology i: an introduction* (2002).
[4] S. H. Strogatz, *Nonlinear dynamics and chaos with student solutions manual: With applications to physics, biology, chemistry, and engineering* (CRC press, 2018).
